# Comparative Evaluation of Commercial Gradient Diffusion and Disk Diffusion Tests for Aztreonam-Avibactam Susceptibility Testing

**DOI:** 10.64898/2026.07.25.740679

**Authors:** Jennifer Pfeiffer, Supram Hosuru Subramanya, Rinki Kumar, Gregory J. Berry, Lars F. Westblade, Daniel A. Green

## Abstract

**Background:** Aztreonam–avibactam (AZA) is a recently approved agent active against metallo-β-lactamase-producing Enterobacterales. However, independent evaluations of commercially available antimicrobial susceptibility testing (AST) products remain limited.

**Methods:** We evaluated 56 carbapenem-resistant Enterobacterales isolates using four newly available AZA AST products, including two gradient diffusion (bioMérieux Etest; Liofilchem MIC Test Strip) and two disk diffusion (Hardy, Liofilchem) tests across three Mueller–Hinton agar manufacturers at two independent clinical laboratories. A diverse isolate collection enriched for nonsusceptible and breakpoint-adjacent MICs was used to rigorously assess categorical agreement. Broth microdilution (BMD) was used as the reference method. Essential agreement (EA), categorical agreement (CA), error rates, and reproducibility were analyzed.

**Results:** All four commercial AST products demonstrated acceptable analytical performance compared with reference BMD. Gradient diffusion demonstrated high EA (bioMérieux 94.3%, Liofilchem 90.5%), whereas CA ranged from 83.3% to 89.0% across all four products. Minor errors accounted for all categorical discrepancies, and occurred almost exclusively among breakpoint-adjacent isolates. Media-related variability was modest, and quality control performance was generally high, with strong within-site precision and consistent performance across laboratories.

**Conclusions:** This study provides the first comprehensive head-to-head comparison of commercial AZA susceptibility testing products and supports their implementation in clinical microbiology laboratories.

## Introduction

Carbapenem-resistant Enterobacterales (CRE) remain a major global public health threat, driven in large part by the increasing prevalence of carbapenemase-producing organisms (1,2). Among these, metallo-β-lactamase (MBL)-producing Enterobacterales present a particular therapeutic challenge, as they hydrolyze nearly all β-lactams and are not inhibited by currently available β-lactamase inhibitors (2,3). As a result, treatment options for infections caused by MBL-producing organisms have historically been limited and often rely on combination regimens with variable efficacy (3,4). Therefore, there is an urgent clinical need for effective therapeutic options supported by reliable susceptibility testing methods.

Aztreonam–avibactam (AZA) is a novel β-lactam/β-lactamase inhibitor combination that restores the activity of aztreonam against organisms co-producing MBLs and serine β-lactamases (3,5–8). Avibactam inhibits class A, C, and some class D β-lactamases, thereby protecting aztreonam from co-produced serine enzymes while aztreonam retains intrinsic stability against hydrolysis by MBLs (3,7,8). This complementary mechanism has led to the development and recent regulatory approval of AZA for the treatment of infections caused by multidrug-resistant Gram-negative organisms, including those harboring MBLs (9).

With the introduction of AZA into clinical practice, clinical microbiology laboratories must establish reliable antimicrobial susceptibility testing (AST) methods to guide therapeutic decision-making. While BMD remains the standard method, routine implementation in clinical laboratories is often limited by workflow, expertise, and staffing constraints. Broth disk elution has also been studied and used, but requires manual and time-consuming setup (10–12). Commercially available gradient diffusion and disk diffusion products therefore represent practical alternatives; however, independent evaluations of their performance for AZA are currently limited, particularly in United States (US) clinical laboratory settings.

A few recent studies have evaluated AZA susceptibility testing using gradient agar or disk diffusion, demonstrating high categorical agreement with reference methods (13–17). However, prior studies have been limited to a single manufacturer’s assay and most have not included a broad enough diversity of isolates to rigorously assess categorical agreement. Furthermore, no previous studies have assessed performance across multiple media manufacturers.

Therefore, in this study, we sought to evaluate and directly compare the performance of four commercially available AZA AST tests, including two gradient diffusion and two disk diffusion platforms, across three Mueller–Hinton agar manufacturers at two independent clinical microbiology laboratories. Using a diverse collection of CRE isolates with well-characterized resistance mechanisms and a wide minimum inhibitory concentration (MIC) distribution, we assessed essential and categorical agreement with reference broth microdilution, as well as analytical precision and inter-laboratory reproducibility. These data provide important insights into the performance characteristics of commercially available AZA testing methods to inform their implementation in clinical laboratories.

## Methods

### Study Design

Fifty-six bacterial isolates obtained from the Centers for Disease Control and Prevention (CDC) Antibiotic Resistance (AR) Isolate Bank and Element Materials Technology (North Liberty, IA) were evaluated to assess the performance of commercially available AZA susceptibility testing products. Two gradient diffusion tests (bioMérieux Etest [bioMérieux, Durham, NC] and Liofilchem MIC Test Strips [Liofilchem, Waltham, MA]) and two disk diffusion products (Hardy Diagnostics [Santa Maria, CA] and Liofilchem disks [Waltham, MA]) were evaluated.

A standardized 0.5 McFarland suspension of each isolate was inoculated onto Mueller-Hinton agar (MHA) from three manufacturers (Remel, Lenexa, Kansas; Hardy; and Becton, Dickinson and Company [BD], Franklin Lakes, NJ). All testing was performed in parallel in the clinical microbiology laboratories at two academic medical centers. Each isolate was evaluated using all four commercial susceptibility testing products on MHA from each of the three manufacturers at both participating laboratories, yielding 336 total observations for each product.

Broth microdilution (BMD) reference MIC values were obtained from CDC and Element Materials Technology and were used as the comparator method. All AST was performed in accordance with Clinical and Laboratory Standards Institute (CLSI) guidelines and manufacturer instructions for use, and interpreted according to US Food and Drug Administration (FDA) breakpoints (18–23). MIC endpoints were read visually (24) according to manufacturer’s instructions (19,22), with all visible growth incorporated into endpoint determination. Any haze, microcolonies, or macrocolonies within 3 mm of the strip were interpreted as growth (19,22). MICs that fell between two doubling dilutions were rounded up to the next doubling dilution. Repeat testing was performed for isolates with major or very major errors, using a fresh subculture and standardized inoculum, with the repeat results used for data analysis. Persistent discrepancies underwent additional testing at an independent reference laboratory, with the reference laboratory MIC used as the final comparator for error classification.

### Bacterial Isolates

A total of 56 Enterobacterales isolates were tested, including *Klebsiella* spp. (n = 21)*, Escherichia coli* (n = 25)*, Enterobacter cloacae* (n = 7)*, Serratia marcescens* (n = 2), and *Proteus mirabilis* (n = 1) (Table 1, Supplemental Table S1). Isolates were selected to represent a broad range of carbapenem resistance mechanisms, including metallo-β-lactamases (MBLs: IMP, NDM, and VIM), serine carbapenemases (KPC and OXA-48-like) and non-carbapenemase-producing CRE. Forty-seven isolates were obtained from the CDC AR Isolate Bank, and nine additional MBL-producing isolates were obtained from Element Materials Technology (Table 1). Reference MICs and FDA interpretations were provided by these sources. The isolate collection was intentionally enriched for isolates with reference MICs spanning the FDA interpretive breakpoints (4-16 µg/ml), with 17 of 56 evaluable isolates (30.4%) falling within this range. Isolates were maintained at −80°C in trypticase soy broth with 20% glycerol. Bacterial isolates were subcultured twice (F1 and F2) on blood agar plates and incubated under suitable growth conditions to obtain fresh, pure colonies. Well-isolated colonies from the overnight F2 culture were then used to prepare the standardized inoculum for susceptibility testing.

**Table 1.**
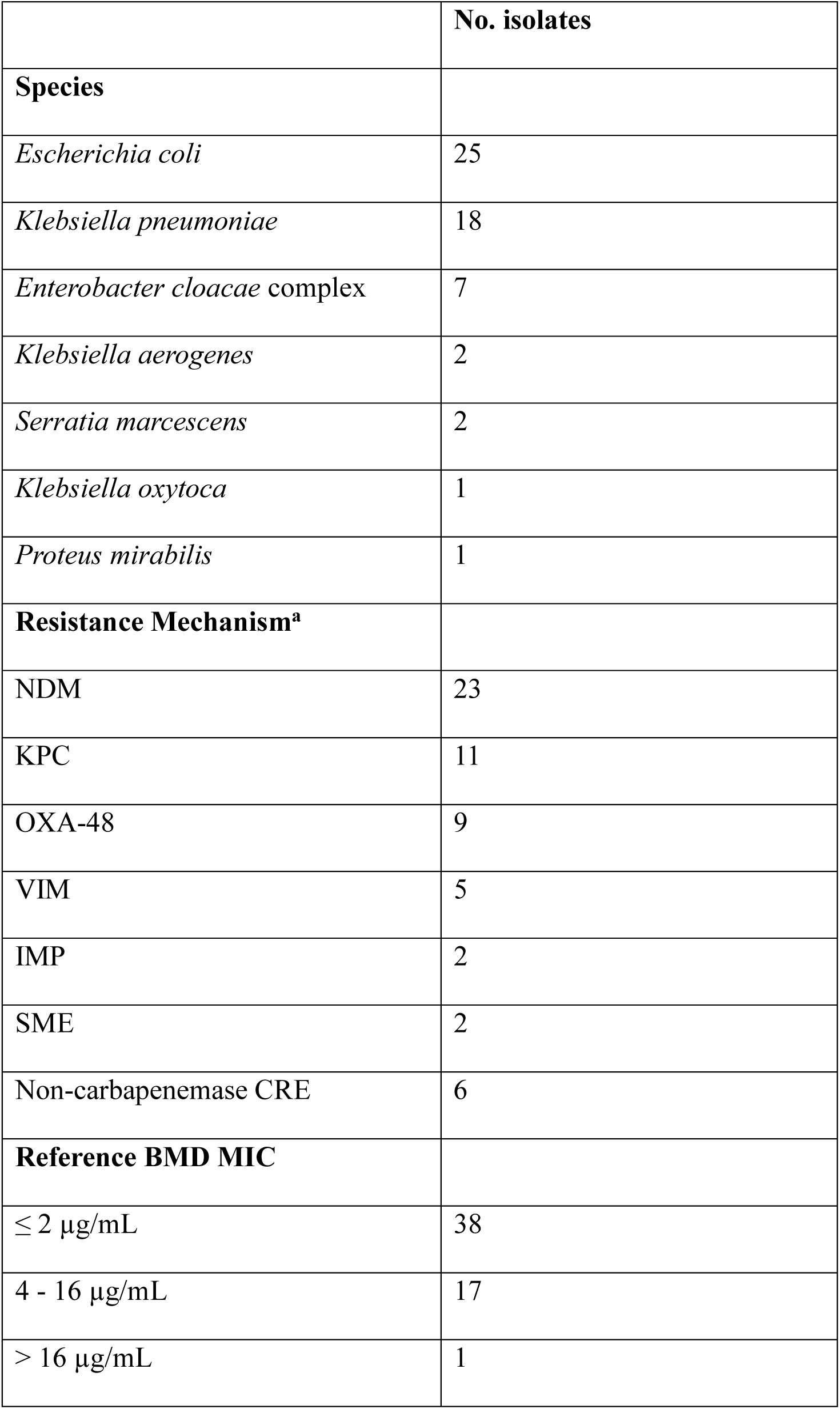
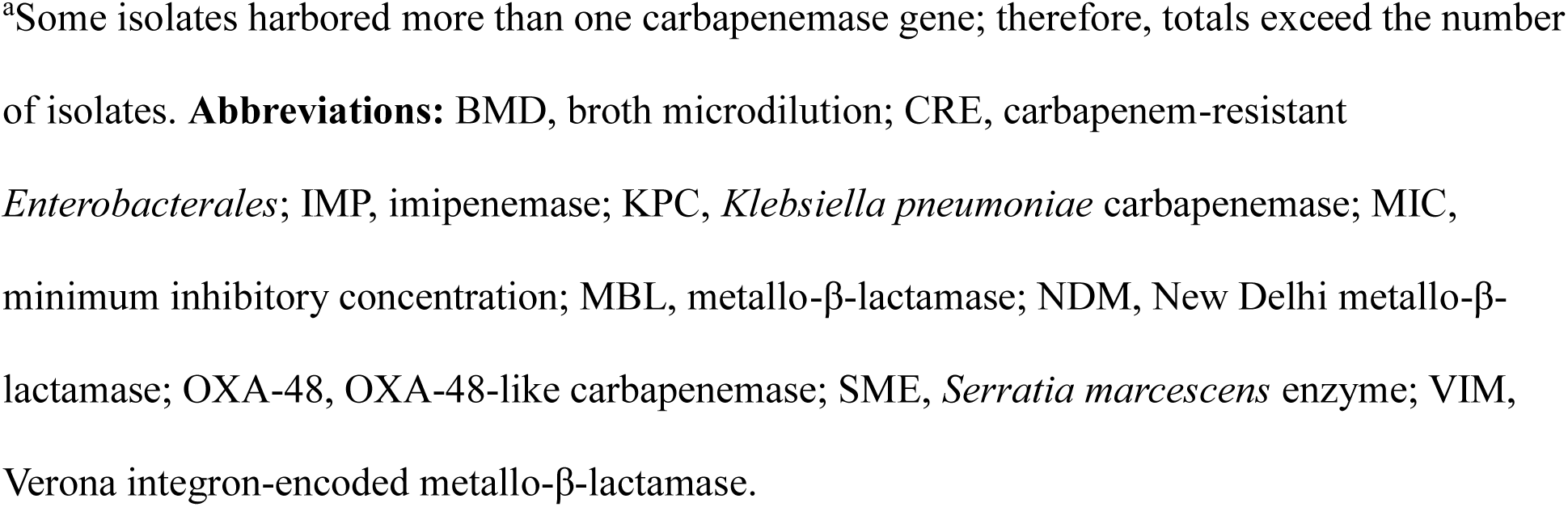
Characteristics of the carbapenem-resistant Enterobacterales isolate collection evaluated in this study.

|  | No. isolates |
| --- | --- |
| **Species** |  |
| *Escherichia coli* | 25 |
| *Klebsiella pneumoniae* | 18 |
| *Enterobacter cloacae* complex | 7 |
| *Klebsiella aerogenes* | 2 |
| *Serratia marcescens* | 2 |
| *Klebsiella oxytoca* | 1 |
| *Proteus mirabilis* | 1 |
| **Resistance Mechanism<sup>a</sup>** |  |
| NDM | 23 |
| KPC | 11 |
| OXA-48 | 9 |
| VIM | 5 |
| IMP | 2 |
| SME | 2 |
| Non-carbapenemase CRE | 6 |
| **Reference BMD MIC** |  |
| ≤ 2 µg/mL | 38 |
| 4 - 16 µg/mL | 17 |
| > 16 µg/mL | 1 |

<sup>a</sup>Some isolates harbored more than one carbapenemase gene; therefore, totals exceed the number of isolates. **Abbreviations:** BMD, broth microdilution; CRE, carbapenem-resistant *Enterobacteriales*; IMP, imipenemase; KPC, *Klebsiella pneumoniae* carbapenemase; MIC, minimum inhibitory concentration; MBL, metallo-β-lactamase; NDM, New Delhi metallo-β-lactamase; OXA-48, OXA-48-like carbapenemase; SME, *Serratia marcescens* enzyme; VIM, Verona integron-encoded metallo-β-lactamase.

### Antimicrobial Susceptibility Testing

Three to five colonies were suspended in sterile saline to achieve a turbidity equivalent to a 0.5 McFarland standard, as determined by visual comparison or by use of an FDA-approved nephelometric device, corresponding to approximately 1.5 × 10^8 CFU/ml. A sterile cotton swab was used to evenly streak the MHA plates to be tested, with the standardized inoculum. AZA 0.016/4-256/4 µg/mL MIC test strips (bioMérieux and Liofilchem) were applied to inoculated MHA plates and incubated for 16-20 hours, according to manufacturer instructions. MICs were recorded at the point of complete inhibition of visible growth. For disk diffusion testing, 30/20 µg AZA disks (Hardy Diagnostics and Liofilchem) were applied to inoculated MHA plates. Plates were incubated for 16-18 hours, according to manufacturer’s instructions. Zone diameters were measured in millimeters and interpreted using FDA breakpoint criteria. All testing was performed on MHA media from three manufacturers (BD, Hardy Diagnostics, and Remel) to assess media-related variability. Results were evaluated individually by site and using combined data from both laboratories.

### Quality Control and Precision

Quality control (QC) testing was performed using American Type Culture Collection (ATCC) strains *E. coli* ATCC 25922, *E. coli* ATCC 35218, and *K. pneumoniae* ATCC 700603. Each QC strain was tested in triplicate on each of five study days using all test products on MHA from each of the three manufacturers at both laboratories using the methods described above. Gradient diffusion MICs and disk diffusion zone diameters were compared against manufacturer-provided QC ranges.

Within-site repeatability for gradient diffusion products was assessed as the proportion of results within ±1 doubling dilution of the site-specific modal MIC for each strain-product combination. Interlaboratory reproducibility was assessed by comparison of site-specific modal MICs and observed MIC ranges. For disk diffusion products, within- and between-laboratory precision was assessed using site-specific mean zone diameters, standard deviations, and observed ranges. QC performance for all products was assessed as the proportion of results within manufacturer-established QC ranges. For QC analysis, MIC values of 0.125 and 0.064 µg/mL were considered equivalent to manufacturer-specified limits of 0.12 and 0.06 µg/mL, respectively.

### Interpretation and Data Analysis

Essential agreement (EA) was defined as the proportion of test MICs within one doubling dilution above or below the reference BMD MIC. Categorical agreement (CA) was defined as the proportion of isolates with matching interpretive categories between test method and reference BMD based on FDA breakpoints: susceptible (≤4 µg/mL), intermediate (8 µg/mL), and resistant (≥16 µg/mL) for gradient diffusion; susceptible (≥ 21 mm), intermediate (18-20 mm), and resistant (≤ 17 mm) for disk diffusion (23).

For both gradient and disk diffusion methods, CA error rates were classified as minor errors, major errors (MEs), or very major errors (VMEs). Minor errors were defined as isolates categorized as intermediate by one method and susceptible or resistant by the comparator method. MEs were defined as isolates categorized as susceptible by BMD but resistant by test method, and VMEs occurred when the reference method categorized an isolate as resistant but the test method categorized it as susceptible.

## Results

### Essential and Categorical Agreement

Overall, all four commercial AST products demonstrated acceptable analytical performance compared with reference BMD, with consistent performance across two laboratories and three Mueller-Hinton agar manufacturers (Table 2, Supplemental Table S2).

**Table 2.**
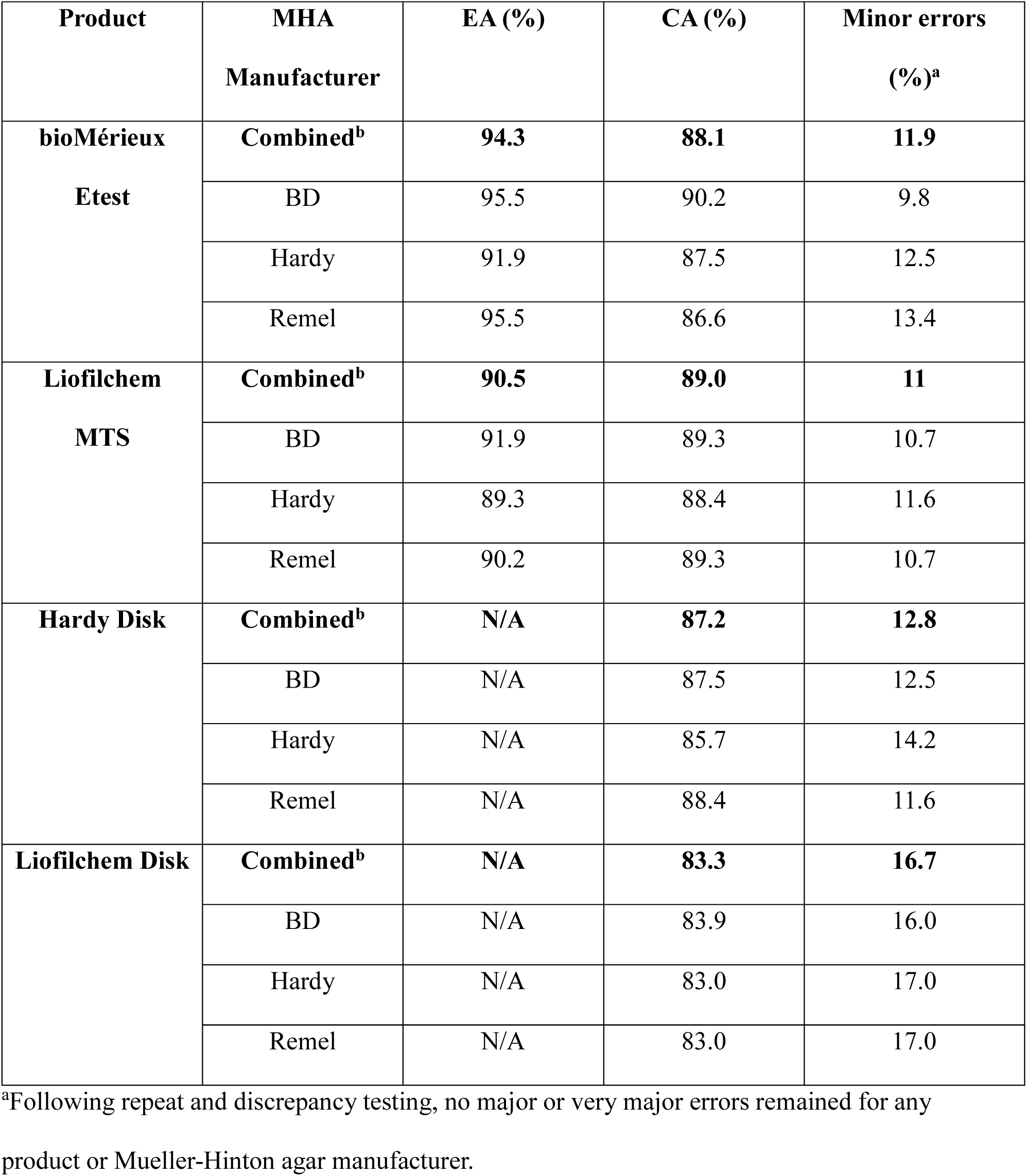
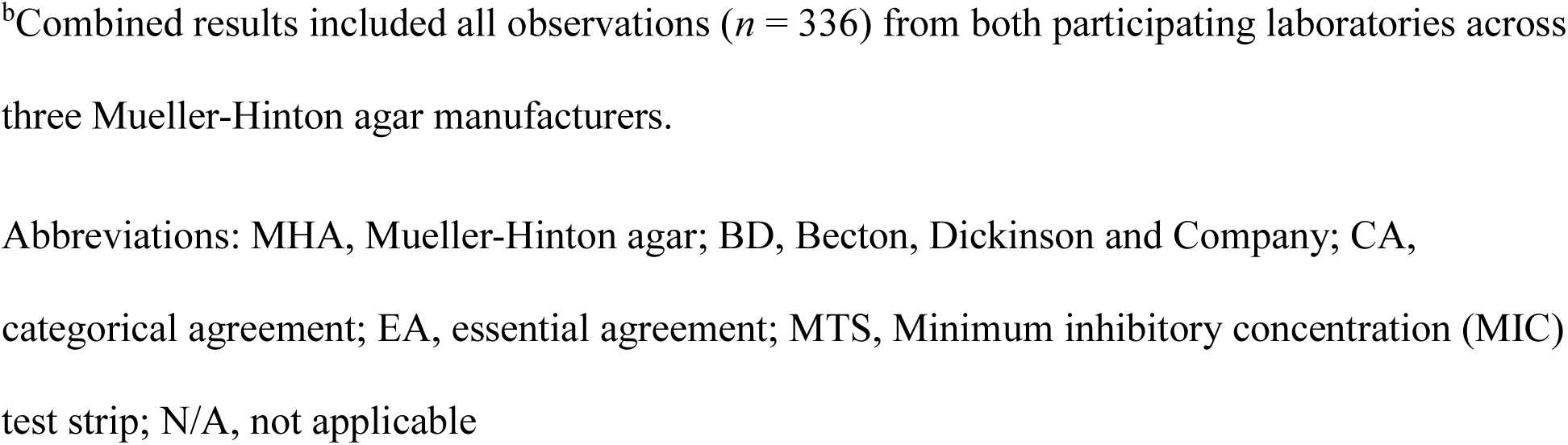
Performance of four commercial aztreonam-avibactam susceptibility testing products compared with reference broth microdilution.

| **Product** | **MHA**<br>**Manufacturer** | **EA (%)** | **CA (%)** | **Minor errors**<br>**(%)<sup>a</sup>** |
| --- | --- | --- | --- | --- |
| **bioMérieux**<br>**Etest** | **Combined<sup>b</sup>** | **94.3** | **88.1** | **11.9** |
|  | BD | 95.5 | 90.2 | 9.8 |
|  | Hardy | 91.9 | 87.5 | 12.5 |
|  | Remel | 95.5 | 86.6 | 13.4 |
| **Liofilchem**<br>**MTS** | **Combined<sup>b</sup>** | **90.5** | **89.0** | **11** |
|  | BD | 91.9 | 89.3 | 10.7 |
|  | Hardy | 89.3 | 88.4 | 11.6 |
|  | Remel | 90.2 | 89.3 | 10.7 |
| **Hardy Disk** | **Combined<sup>b</sup>** | **N/A** | **87.2** | **12.8** |
|  | BD | N/A | 87.5 | 12.5 |
|  | Hardy | N/A | 85.7 | 14.2 |
|  | Remel | N/A | 88.4 | 11.6 |
| **Liofilchem Disk** | **Combined<sup>b</sup>** | **N/A** | **83.3** | **16.7** |
|  | BD | N/A | 83.9 | 16.0 |
|  | Hardy | N/A | 83.0 | 17.0 |
|  | Remel | N/A | 83.0 | 17.0 |

<sup>a</sup>Following repeat and discrepancy testing, no major or very major errors remained for any product or Mueller-Hinton agar manufacturer.
<sup>b</sup>Combined results included all observations ( $n = 336$ ) from both participating laboratories across three Mueller-Hinton agar manufacturers.
Abbreviations: MHA, Mueller-Hinton agar; BD, Becton, Dickinson and Company; CA, categorical agreement; EA, essential agreement; MTS, Minimum inhibitory concentration (MIC) test strip; N/A, not applicable

For gradient diffusion, bioMérieux Etest demonstrated slightly higher overall EA (94.3%) than Liofilchem MIC Test Strip (90.5%), while overall CA was similar for both products (88.1% and 89.0%, respectively). Overall CA for disk diffusion was 83.3% - 87.2%, with Hardy disks demonstrating slightly higher agreement than Liofilchem. Following repeat and discrepancy testing, there were no major or very major errors for any of the gradient or disk diffusion tests, and all remaining categorical discrepancies were minor errors. Results of the initial, repeat, and discrepancy testing for isolates with initial major or very major errors are shown in Supplemental Table S3.

Performance was similar between the two participating laboratories (Supplemental Table S2). Overall EA, CA, and error rates differed by only a few percentage points between laboratories for all four products, and the relative performance of each method was consistent across sites.

### Error Analysis

Categorical errors occurred almost exclusively among isolates with reference MICs near the FDA interpretive breakpoints (4-16 ug/ml), where small differences in MIC determination can result in categorical shifts (25,26). Following repeat and discrepancy testing, all remaining discrepancies were minor errors. Importantly, 96.1% of minor errors with gradient diffusion products were within EA, highlighting that these errors were driven by single doubling-dilution differences in MIC determination rather than substantial analytical disagreement.

MIC distribution analysis demonstrated that most gradient diffusion results clustered on or immediately adjacent to the line of absolute agreement with reference BMD (Figures 1 and 2). No consistent directional bias in MIC determination was observed for either gradient diffusion product.

**Figure 1.**
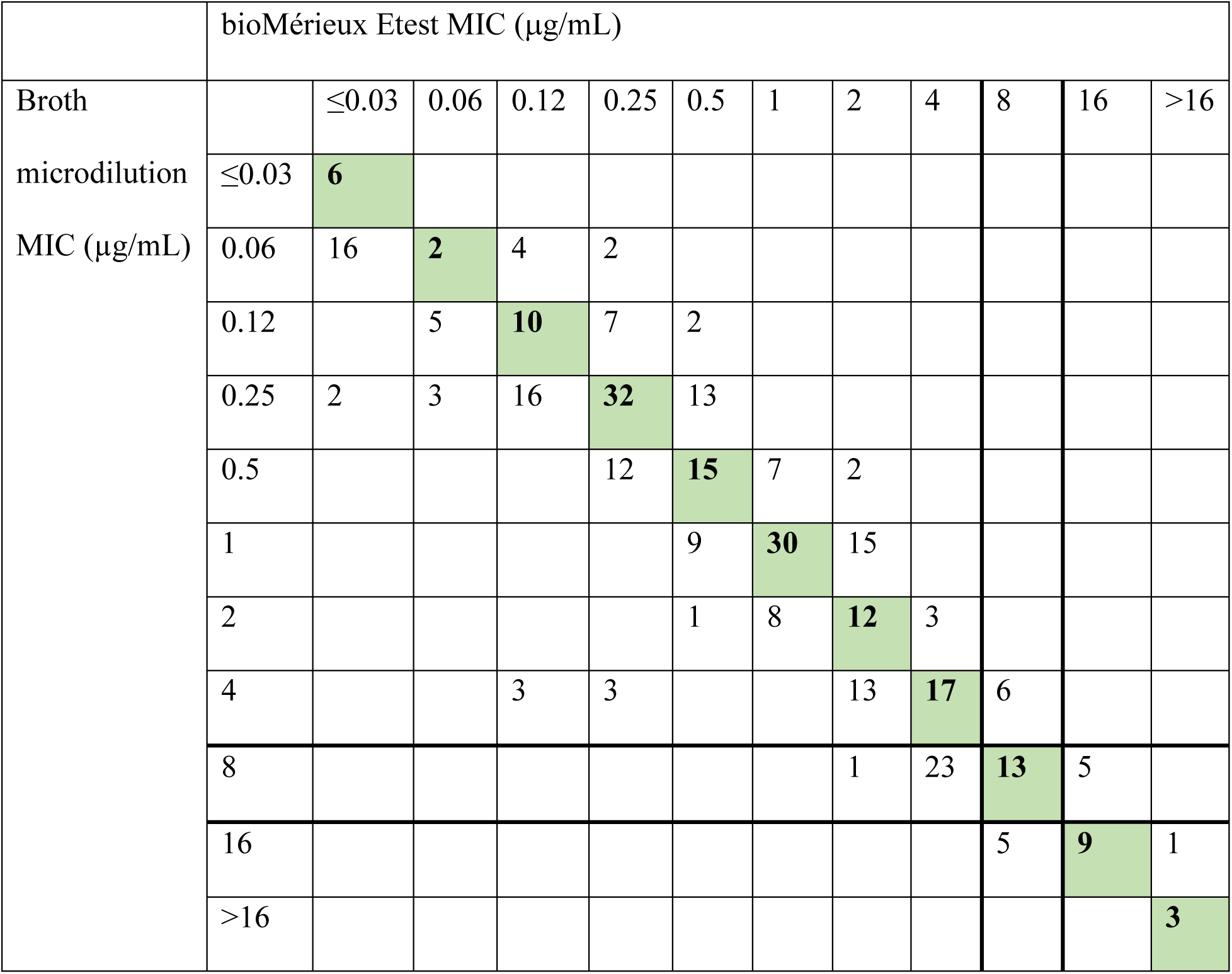
Comparison of bioMérieux Etest aztreonam-avibactam MICs with reference broth microdilution MICs. Cell values indicate the number of observations among 336 determinations obtained across three Mueller-Hinton agar manufacturers and two laboratories. Diagonal shaded cells represent absolute agreement; cells immediately adjacent to the diagonal represent essential agreement. Bold lines delineate FDA interpretive breakpoints.

**Figure 2.**
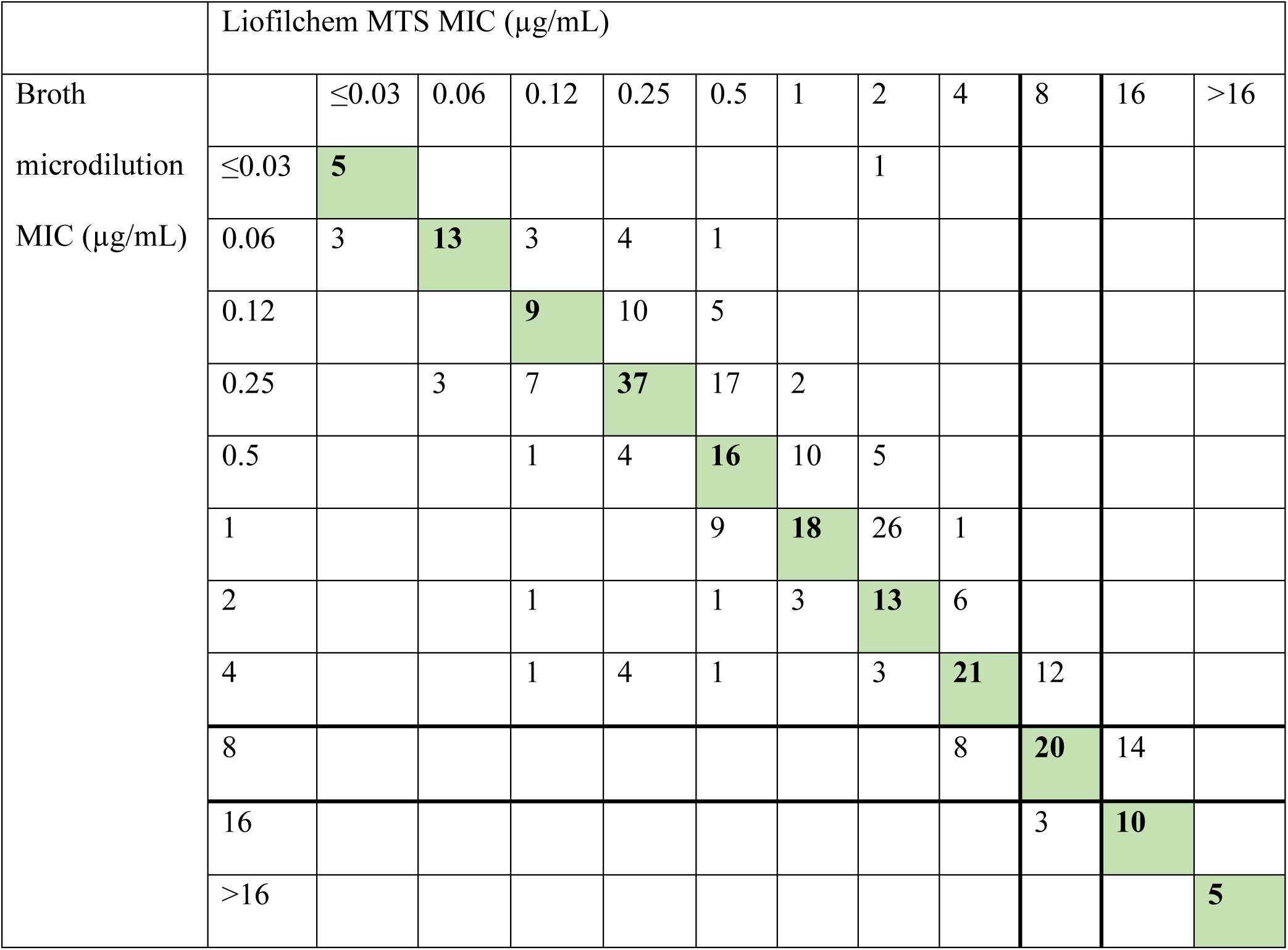
Comparison of Liofilchem MIC Test Strip aztreonam-avibactam MICs with reference broth microdilution MICs. Cell values indicate the number of observations among 336 determinations obtained across three Mueller-Hinton agar manufacturers and two laboratories. Diagonal shaded cells represent absolute agreement; cells immediately adjacent to the diagonal represent essential agreement. Bold lines delineate FDA interpretive breakpoints.

### Media-Related Variability

Only modest differences in performance were observed among the three Mueller-Hinton agar manufacturers (Table 2), and no single Mueller-Hinton agar manufacturer consistently demonstrated superior performance across all products.

For bioMérieux Etest, EA and CA ranged from 92.0% - 95.5% and 84.5% - 90.2%, respectively, while EA and CA for Liofilchem MIC Test Strips ranged from 89.3% - 92.0% and 88.4% - 89.3%, respectively. BD and Remel media yielded slightly higher EA than Hardy media for both gradient diffusion products, while differences in CA were smaller and less consistent. Disk diffusion performance varied little by media, with CA differing by no more than 3% across the three agar manufacturers for either disk product.

### Quality control and Precision

QC performance and analytical precision are summarized in Table 3. All bioMérieux Etest QC results were within manufacturer-established ranges. For the Liofilchem MIC Test Strip, 93.3%, 60.0%, and 100% of results were within the manufacturer-established QC ranges for *E. coli* ATCC 25922, *E. coli* ATCC 35218, and *K. pneumoniae* ATCC 700603, respectively. Within-site repeatability was high for both gradient diffusion products, with 100% of results within ±1 doubling dilution of the site-specific modal MIC for every strain-product combination at both laboratories.

**Table 3.** Quality control performance and precision of four commercial aztreonam-avibactam susceptibility testing products.

A. Gradient Diffusion
| Product | QC strain | Manufacturer QC range (µg/mL) | Within manufacturer QC range, n/N (%) | Site 1: modal MIC (range), µg/mL | Site 2: modal MIC (range), µg/mL | Within ±1 DD of site-specific modal MIC, Site 1 / Site 2, n/N (%) |
| --- | --- | --- | --- | --- | --- | --- |
| bioMérieux Etest | *E. coli* ATCC 25922 | 0.032–0.125 | 90/90 (100) | 0.064 (0.032–0.125) | 0.064 (0.032–0.125) | 45/45 (100) / 45/45 (100) |
| bioMérieux Etest | *K. pneumoniae* ATCC 700603 | 0.064–0.5 | 90/90 (100) | 0.25 (0.125–0.25) | 0.25 (0.125–0.25) | 45/45 (100) / 45/45 (100) |
| Liofilchem MTS | *E. coli* ATCC 25922 | 0.03–0.12 | 84/90 (93.3) | 0.125 (0.064–0.25) | 0.125 (0.125–0.25) | 45/45 (100) / 45/45 (100) |
| Liofilchem MTS | *E. coli* ATCC 35218 | 0.016–0.06 | 54/90 (60.0) | 0.125 (0.064–0.25) | 0.064 (0.032–0.064) | 45/45 (100) / 45/45 (100) |
| Liofilchem MTS | *K. pneumoniae* ATCC 700603 | 0.06–0.5 | 90/90 (100) | 0.25 (0.125–0.5) | 0.25 (0.25–0.5) | 45/45 (100) / 45/45 (100) |

| Product | QC strain | Manufacturer<br>QC range<br>(mm) | Within<br>manufacturer<br>QC range,<br>n/N (%) | Site 1: mean ±<br>SD (range),<br>mm | Site 2: mean<br>± SD<br>(range), mm |
| --- | --- | --- | --- | --- | --- |
| Hardy disk | *E. coli*<br>ATCC<br>25922 | 32–38 | 89/90 (98.9) | 33.3 ± 1.2 (31–<br>37) | 34.1 ± 1.7<br>(32–38) |
| Hardy disk | *E. coli*<br>ATCC<br>35218 | 31–38 | 90/90 (100) | 32.9 ± 1.1 (31–<br>35) | 34.6 ± 1.6<br>(32–38) |
| Hardy disk | *K. pneumoniae*<br>ATCC<br>700603 | 26–32 | 90/90 (100) | 28.2 ± 1.0 (26–<br>31) | 28.0 ± 1.5<br>(26–30) |
| Liofilchem<br>disk | *E. coli*<br>ATCC<br>25922 | 32–38 | 88/90 (97.8) | 34.5 ± 1.5 (30–<br>38) | 35.7 ± 1.3<br>(34–38) |
| Liofilchem<br>disk | *E. coli*<br>ATCC<br>35218 | 31–38 | 90/90 (100) | 34.1 ± 1.2 (32–<br>37) | 35.8 ± 1.2<br>(34–38) |
| Liofilchem<br>disk | *K. pneumoniae*<br>ATCC<br>700603 | 26–32 | 90/90 (100) | 30.0 ± 1.0 (28–<br>32) | 29.1 ± 1.5<br>(27–31) |

QC testing was performed in triplicate on five study days using Mueller-Hinton agar from three manufacturers at two independent laboratories (90 observations per product–QC strain combination; 45 observations per laboratory). QC performance was assessed using manufacturer- established QC ranges. For gradient diffusion methods, site-specific modal MICs and observed ranges are shown, and within-site repeatability was assessed as the proportion of results within $\pm 1$ doubling dilution (DD) of the site-specific modal MIC. For disk diffusion methods, site- specific results are presented as mean ± standard deviation (SD) and observed range. MIC values of 0.125 and 0.064 $\mu\text{g/mL}$ were considered equivalent to manufacturer-specified QC limits of 0.12 and 0.06 $\mu\text{g/mL}$ , respectively.

The most notable QC excursion occurred with *E. coli* ATCC 35218 tested using the Liofilchem MIC Test Strip. The modal MIC was 0.125 µg/mL at Site 1 compared with 0.064 µg/mL at Site 2. At Site 1, minimal growth extended to the 0.125 µg/mL endpoint, resulting in MICs above the manufacturer-established upper QC limit of 0.06 µg/mL. This pattern was reproducible across all five testing days at Site 1, was not observed at Site 2, and was limited to this strain-product combination.

Disk diffusion demonstrated excellent QC compliance, with 98.9–100% of Hardy disk and 97.8–100% of Liofilchem disk results falling within manufacturer-established QC ranges. Zone diameter distributions were generally similar between laboratories, with site-specific means differing by ≤1.7 mm across all strain-product combinations (Table 3).

## Discussion

The recent regulatory approval of AZA provides an important therapeutic option for infections caused by CRE, particularly those harboring MBLs, for which treatment options have historically been limited. As AZA enters routine clinical practice, clinical microbiology laboratories require practical, standardized susceptibility testing to guide its therapeutic use. This study provides the most comprehensive comparative evaluation of commercially available susceptibility testing products for AZA to date, evaluating and directly comparing four agar-based tests across three MHA manufacturers at two independent laboratories. Overall, both gradient and disk diffusion products demonstrated acceptable analytical performance compared with reference BMD, supporting their implementation into routine clinical laboratory testing.

Across both laboratories and all media types, both gradient diffusion products showed high EA, whereas CA was more moderate across all four products. These differences highlight an important consideration when evaluating new AST methods. EA primarily reflects the analytical performance of the assay, whereas CA can be strongly influenced by the MIC distribution of study isolates. By deliberately enriching our isolate set with breakpoint-adjacent organisms, we created a more rigorous assessment of CA than most previous evaluations. Under these conditions, expected analytical variability frequently resulted in categorical discrepancies despite remaining within EA. Reassuringly, there were no major or very major errors after repeat and discrepancy testing, and 96.1% of minor errors by gradient diffusion were within EA. Two isolates that initially produced VMEs with at least one commercial product underwent discrepancy testing at an independent laboratory and were subsequently reclassified as minor errors. Collectively, these findings support the acceptable analytical performance of all four products under a deliberately challenging study design while emphasizing that CA should be interpreted in the context of the MIC distribution of the isolates evaluated.

An important strength of this study was a thorough assessment of analytical precision across two laboratories and three MHA manufacturers. Overall precision was high, although a reproducible, site-specific upward MIC shift was observed for *E. coli* ATCC 35218 with the Liofilchem MIC Test Strip, resulting in reduced QC compliance. This shift was associated with minimal trailing growth at Site 1 and was limited to this strain-product combination, suggesting a systematic testing effect rather than generalized imprecision. Disk diffusion products likewise demonstrated high QC compliance and similar zone diameter distributions between laboratories.

Our study findings build upon a growing body of literature supporting the performance of agar-based AZA susceptibility testing products. Chan *et al.* compared Liofilchem gradient and disk diffusion products with broth disk elution on 31 CRE isolates and observed 100% CA (13). Lemon *et al.* likewise reported high CA (98.5%) for Liofilchem strips using a mix of clinical isolates and isolates from the CDC AR Bank (16). Both studies included relatively few breakpoint-adjacent organisms, which limited the opportunity to observe categorical discrepancies driven by single dilution MIC variation. Hamze *et al.* also reported high CA for Liofilchem gradient strips (97.2%), but did not specify the MIC distribution of the study isolates (15). By contrast, Deschamps *et al.* included more breakpoint-adjacent isolates and reported findings similar to ours; Liofilchem strip EA was 91.7% and CA was 89.7%, with 10.3% minor errors (14). More recently, Bui *et al.* evaluated bioMérieux Etest in a multicenter study of over 600 clinical and challenge Enterobacterales isolates and observed excellent performance (95.7% EA and 98.0% CA using FDA breakpoints), although relatively few isolates had MICs near the FDA interpretive breakpoints (17).

Collectively, these studies support the utility of agar-based AZA testing products, while our study expands on these analyses in several important ways. Unlike previous studies that evaluated a single product, we directly compared four commercial assays, including both gradient diffusion and disk diffusion methods. To our knowledge, this is the first study to evaluate Hardy disk diffusion products and the first to systematically assess media-related variability. In addition, our isolate set was enriched for breakpoint-adjacent MICs, providing a more rigorous assessment of CA than most prior studies. Taken together, this study provides the first comprehensive head-to-head comparison of commercially available AZA susceptibility testing products across multiple laboratories, MHA manufacturers, and a deliberately challenging isolate collection.

Prior to the availability of commercial assays and broth disk elution for AZA, clinical laboratories relied on various methods of synergy testing to derive MICs for the combination of ceftazidime/avibactam and aztreonam (25,26). Synergy methods are significantly more time-consuming and labor-intensive, while also lacking methodological standardization, subjecting them to interpretive challenges. The FDA clearance and commercial availability of multiple disk diffusion and gradient agar diffusion assays provide a more convenient and accessible alternative that is easier for laboratories to validate and incorporate into routine clinical testing. Our findings, together with those of previous studies, support the transition from synergy testing methods to these commercially available assays for routine AZA testing.

This study has several limitations. The isolate collection was moderate in size, though each isolate was evaluated using four commercial susceptibility testing products across three MHA manufacturers at two independent laboratories, resulting in 336 observations for each product. The isolate collection was also enriched for carbapenemase-producing *Enterobacterales,* especially isolates with MICs spanning the FDA interpretive breakpoints. While this study design provided a rigorous assessment of CA, it likely produced a more conservative estimate of CA than would be expected during routine clinical testing, where the majority of isolates have MICs well removed from the interpretive breakpoints. Intermediate and resistant isolates remain uncommon, and currently available panels from CDC and other repositories lack extensive collections of non-susceptible strains. As additional isolates become available, future studies will be able to more comprehensively evaluate the performance of these products in non-susceptible organisms. Our evaluation was also limited to CRE, reflecting the principal intended clinical use for AZA, though future studies are also needed to understand the performance characteristics for other organisms, such as *Pseudomonas* and *Stenotrophomonas.* Additionally, a few of the tested isolates demonstrated growth patterns that complicated MIC or zone size interpretation, reflecting challenges that may also be encountered in routine clinical laboratory practice. Finally, although our study was performed at two independent laboratories, additional multicenter evaluations would provide further data on the reproducibility and generalizability of these products across a wider range of laboratory environments.

In summary, commercially available disk diffusion and gradient diffusion products for AZA susceptibility testing demonstrated reliable analytical performance compared with BMD, with high analytical precision across media types and laboratories. While minor discrepancies were observed among breakpoint-adjacent isolates, these findings reflect expected analytical variability rather than intrinsic limitations of the products. Our results support the implementation of these products in clinical laboratories and provide practical guidance for AZA susceptibility testing as its use expands.

## Acknowledgements

The authors thank Michael Hoy (AbbVie) for his assistance in identifying appropriate CDC Antibiotic Resistance Isolate Bank panels to support assembly of the study isolate collection.

